# Growth of tumor emboli within a vessel model reveals dependence on the magnitude of mechanical constraint

**DOI:** 10.1101/2020.07.06.190447

**Authors:** Jonathan Kulwatno, Jamie Gearhart, Xiangyu Gong, Nora Herzog, Matthew Getzin, Mihaela Skobe, Kristen L. Mills

## Abstract

Tumor emboli – aggregates of tumor cell within vessels – pose a clinical challenge as they are associated with increased metastasis and tumor recurrence. When growing within a vessel, tumor emboli are subject to a unique mechanical constraint provided by the tubular geometry of the vessel. Current models of tumor emboli use unconstrained multicellular tumor spheroids, which neglect this mechanical interplay. Here, we modelled a lymphatic vessel as a 200 μm-diameter channel in either a stiff or soft, bioinert agarose matrix, and we modelled colon or breast cancer tumor emboli with aggregates of HCT116 or SUM149PT cells, respectively. The stiff vessel model constrained the tumor emboli to the cylindrical geometry, which led to continuous growth of the emboli, in contrast to the growth plateau that unconstrained spheroids exhibit. Emboli morphology in the soft vessel model, however, was dependent on the magnitude of mechanical mismatch between the vessel matrix and the cell aggregates. In general, when the elastic modulus of the vessel was greater than the emboli (*E_ves_* / *E_emb_* > 1), the emboli were constrained to grow within the vessel geometry, and when the elastic modulus of the vessel was less than the emboli (0 < *E_ves_* / *E_emb_* < 1), the emboli bulged into the matrix. Inhibitors of myosin-related force generation decreased the elastic modulus and/or increased the stress relaxation of the tumor cell aggregates, effectively increasing the mechanical mismatch. The increased mechanical mismatch after drug treatment was correlated with increased confinement of tumor emboli growth along the vessel, which may translate to increased tumor burden due to the increased tumor volume within the diffusion distance of nutrients and oxygen.

**INSIGHT BOX:** The growth of tumor emboli—aggregates of tumor cells within vessels—is associated with aggressive cancer progression and metastasis. Models of their growth have not taken into account their biomechanical context, where radial expansion is constrained, but lengthwise expansion is free in the vessel. Here, we modelled the vessel geometry with a cylindrical microchannel in a hydrogel. In contrast to unconstrained or fully embedded aggregates, vessel-like constraint promotes growth of emboli in our model. The growth advantage is increased when the matrix is stiffened or actomyosin contractility weakened, both of which effectively increase the magnitude of mechanical constraint. This study sheds light on increased tumor burden in vessel-based growth and indicates a need to study tumor progression in similar environments.

## INTRODUCTION

Presence of tumor emboli in the lymphatic and blood vasculature is associated with increased metastasis and tumor recurrence and has been correlated with poor patient prognosis [1–4]. Tumor emboli occur when, as part of the metastatic cascade, tumor cells invade into vasculature and grow within the vessels. Lymphatic vessels, in particular, frequently contain tumor emboli in many cancer types, including breast and colorectal cancer, before lymph node metastases can be detected. In addition to transporting tumor cells to the lymph node, lymphatics are also niches for growth of intravascular tumor lesions initiated by arrested tumor emboli, a type of metastatic growth referred to as in transit metastases [5–9]. Despite their importance for metastatic spread, little is understood about what factors influence the growth of tumor emboli [4,10–12]. Particularly, it is not yet understood how the biomechanical properties of the emboli and of the surrounding extracellular microenvironment may affect their growth. Appropriate *in vitro* models of the specific environment of tumor emboli, taking into consideration factors such as geometry and matrix mechanical properties, are necessary to study such interactions.

Currently, the progression of tumor emboli is studied *in vitro* using multicellular tumor spheroids (MCTSs) [10,13,14]. Most MCTS models derive from the aggregation of tumor cells on non-adhesive surfaces. MCTSs are three-dimensional structures that better recapitulate *in vivo* tumors in form and behavioral responses than 2D Petri dish culture [15–17]. However, they only account for tumor cell-tumor cell interactions and neglect interactions with their microenvironments. Solid tumors grow under mechanical constraints provided by their surrounding tissue [18–20]. As such, there is an interplay between a tumor and its microenvironment due to the competition for space: a growing tumor expands against the surrounding tissue while the surrounding tissue confines the tumor. Experimentalists have isolated this biomechanical crosstalk by fully embedding tumor cells within tissue-mechanics-mimicking hydrogels, which has provided insights into how growth and disease progression are influenced by the compressive stress provided by the microenvironment [18–25]. Compressive stress has been shown to inhibit growth of tumor cells and tumor cell aggregates by affecting proliferation and apoptosis, but also stimulating the migration of carcinoma cells [19–21,24].

Studies of compressive stress on tumors have focused mainly on uniform stresses acting on solid tumors, assuming that the solid tumor is confined within the connective tissue. However, in the context of tumor emboli, the tumor is only constrained by the geometry of the vessel, which can be simplified as a cylindrical channel. Here, a growing tumor embolus freely expands until it reaches the boundaries of the vessel, at which point continued expansion is constrained only at the circumference of the vessel to a degree proportional to the stiffness of the surrounding tissue. Therefore, the model microenvironment of a tumor emboli—a vessel-like constraint model—lies between that of no constraint, as the MCTS models, and fully embedding tumor cells or tumor cell aggregates within a hydrogel matrix. To our knowledge, a vessel-like constraint model has not yet been used to study the mechanics of tumor emboli growth.

In this study, we aimed to gain insights into how biomechanical interactions between tumor emboli and the extracellular matrix influence the growth of tumor emboli within the lymphatic vasculature. We modeled a lymphatic vessel as a cylindrical microchannel formed within a tissue-mechanics-mimicking hydrogel and compared the growth and morphology of unconstrained MCTSs to those constrained by our model vessel-like constraint. Changing the mechanical properties of the vessel-like constraint modified the growth and morphology of model tumor emboli from two cell lines: human colon cancer cells (HCT116) and human inflammatory breast cancer cells (SUM149PT) that represent cells derived from cancers where tumor emboli are frequently observed [4,26,27]. Further, the growth behavior—whether the model emboli were confined to the channel geometry or bulged into the matrix—was determined by the mismatch in the mechanical properties between the hydrogel matrix and tumor cell aggregates, which was modulated by inhibition of myosin-related force generating mechanisms.

## MATERIALS & METHODS

### Cell culture

HCT116 human colon cancer cells were purchased from ATCC. Cells were grown as monolayers in McCoy’s 5A medium (Sigma) supplemented with 10% FBS (Gibco) and 1x antimycotic/antibiotic solution (Gibco) in an incubator at 37 °C and 5% CO_2_. Upon reaching 80-90% confluency, cells were collected using 0.05% trypsin (Gibco), centrifuged to remove the collection media, and replaced with fresh media.

SUM149PT human inflammatory breast cancer cells were generously provided by Dr. Jason Herschkowitz of the University at Albany. Cells were grown as monolayers in Ham’s F-12 medium (Biowhitaker) supplemented with 5% FBS (Gibco), 1x antimycotic/antibiotic solution (Gibco), 5 μg/mL insulin (Gibco), and 1 μg/mL hydrocortisone (Acros) in an incubator at 37 °C and 5% CO_2_. Upon reaching 80-90% confluency, cells were collected using 0.05% trypsin (Gibco), centrifuged to remove the collection media, and replaced with fresh media.

### Tumor spheroid generation and growth

In a 96-well plate, 50 μL of molten 1.0% agarose was pipetted into each well and placed at 4 °C for 30 minutes to gel. After gelation, 200 μL of the respective media for each cell line was pipetted into the wells. Cells were then seeded at 1000 cells per well and centrifuged for 10 minutes at 200 × g. The spheroids were cultured in an incubator at 37 °C and 5% CO_2_ for 20 days for growth studies, 10 days for mechanical testing and drug inhibition experiments, and 7 days for western blotting. Brightfield images of the tumor spheroids were taken daily using an Axio Vert.A1 (Zeiss).

### Template for molding cylindrical channels in hydrogels

Three-dimensional printed templates were used to aid in the fabrication of cylindrical channels in agarose hydrogels. The templates were printed with acrylonitrile butadiene styrene (ABS) using a UPrint SE Plus (Stratasys). The templates consisted of a walled rectangular opening with inlets on opposite ends to accommodate a trimmed needle guide on either end. A solid rod—200 μm diameter (Seirin)—was threaded through the two needles to span the rectangular opening of the template (Figure 1A). Polydimethylsiloxane (PDMS) was poured into two large U-shaped pockets wrapped around either side of the rectangular opening and cured. A trimmed microscope slide (Globe Scientific) was placed below the template and secured by binder clips that bonded the slide to the PDMS, ensuring a seal between the glass and the template. All components were either soaked in 70% ethanol overnight or kept under UV for sterilization prior to assembly. The needles were coated with sterile silicone oil during assembly to decrease agarose adhesion.

**Figure 1.**
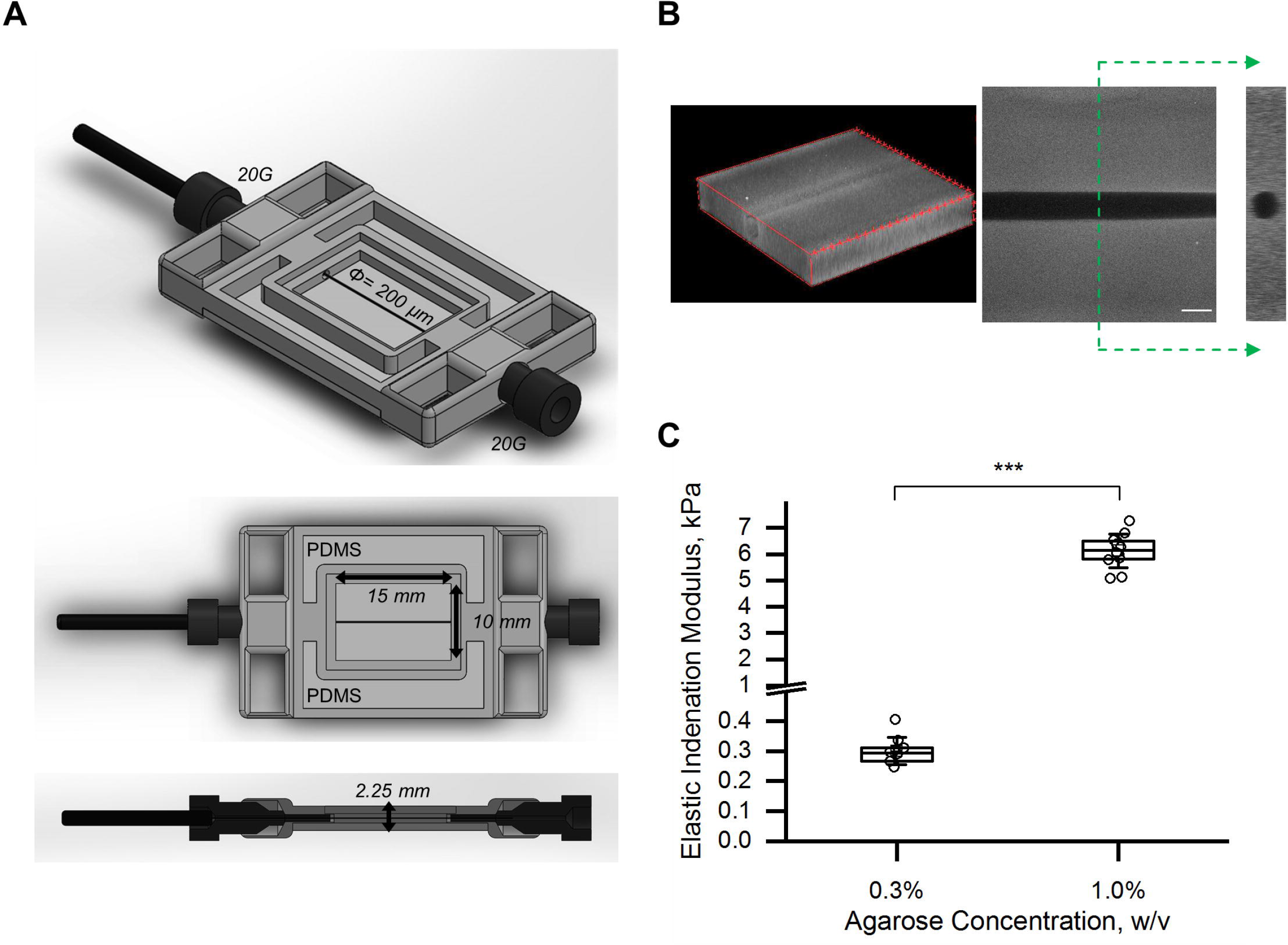
Microchannels fabricated as models to provide a vessel-like constraint. A) The shorter 3D printed template used to aid in the fabrication of cylindrical microchannels in agarose hydrogels. A 200 μm-diameter rod was suspended across the rectangular opening into which agarose was dispensed and gelled. Removal of the rod formed the vessel-like model. B) A 3D reconstruction and orthographic projections (top and side) of the vessel-like model. Fluorescent microbeads were mixed into the agarose prior to gelation to confirm the integrity of the microchannel. Scale bar is 200 μm. C) Elastic indentation moduli of agarose gels at different concentrations. Data is representative of n ≥ 3 from three independent experiments.

The suspended 200 μm-diameter rod was used as a template to mold cylindrical channels in agarose hydrogels (Sigma, low-gelling temperature). Two different agarose concentrations were used: 0.3% (w/v) to provide a *soft* matrix and 1.0% (w/v) to provide a *stiff* matrix. In either case, agarose powder was dissolved in PBS at 100 °C then cooled to 40 °C before use. The templates were pre-cooled at 4 °C for 10 minutes prior to addition of molten agarose, which was pipetted into the rectangular opening of assembled templates. The agarose was allowed to cool at 4 °C for 20 minutes, after which the solid rod and the needles were carefully removed. To confirm the integrity of the microchannel, fluorescent microbeads of 200 nm (ThermoFisher) were mixed in an 0.3% agarose solution at a concentration of 5.30 × 10^9^ beads/mL before gelation and then imaged (Figure 1B).

Two different templates were produced that only differed in the dimension of the rectangular opening in order to accommodate higher aspect ratio growth of tumor cells in the *stiff* agarose. In particular, the shorter template had inner dimensions of 15 mm (length) × 10 mm (width) × 2.25 mm (depth), which held 450 μL of *soft*, 0.3% (w/v), agarose. The inlets on opposite ends of the rectangular opening were 0.908 mm in diameter (Figure 1A) through which two 20-gauge needles (20G, O.D. = 0.908 mm; B.D.) were inserted. The longer template had inner dimensions of 30 mm (length) × 8.5 mm (width) × 1.75 mm (depth) (Supplementary Figure S1), which held 750 μL of the *stiff*, 1.0% (w/v), agarose. The inlets on opposite ends of the rectangular opening were 0.819 mm in diameter (Supplementary Figure S1) through which a normal and a trimmed 21-gauge needle (21G, O.D. = 0.819 mm; B.D.) were inserted.

### Tumor emboli generation and growth

A cell suspension containing either HCT116 cells at 1×10^5^ cells/mL or SUM149PT cells at 3×10^5^ cells/mL was injected with a pipette into the channel through the inlets. Flow through the now-hollow microchannel was aided by brief and shallow tilting of the template. The microchannel-containing hydrogels were then gently transferred out of the templates and into Petri dishes and incubated while the cells were allowed to form aggregates within the microchannels. Once the aggregates reached the diameter of the microchannel, which took between 5 and 7 days, they were deemed emboli. Emboli-containing samples were cultured in clean Petri dishes (to eliminate the cells that had leaked out of the channel and attached to the dish) in an incubator for 20 days for growth studies or 10 days for drug inhibition experiments. Brightfield images of the tumor emboli were taken daily using an Axio Vert.A1 (Zeiss).

### Micro computed tomography (microCT)

To examine the three-dimensional morphology of tumor emboli grown in both stiff and soft matrices, we imaged samples using microcomputed tomography (microCT). Samples were incubated in an iodine solution overnight (Acros). A microCT scan was acquired using a vivaCT 40 (Scanco Medical) at 45 kVp and 177 μA with a pixel size of 10.5 μm × 10.5 μm × 10.5 μm. IMARIS software (Oxford Instruments) was used to reconstruct the scan.

### Inhibition of force generation via drugs

To examine the effects of inhibiting force generation, we used the ROCK inhibitor Y-27632 (Enzo Life Sciences) at 100 μM and the myosin II inhibitor blebbistatin (Enzo Life Sciences) at 10 μM. For unconstrained samples, spheroids were grown for three days to allow compaction and then the media was replaced with media containing a drug, the solvent of a drug (water (H_2_O) for Y-27632 or dimethyl sulfoxide (DMSO) for blebbistatin), or no drug (normal). Samples were then cultured for seven more days before being mechanically tested. Similarly, for microchannel-constrained samples, cells were grown in the channels until they reached the diameter of the microchannel and then the emboli-containing samples were transferred into 6-well plates with media containing a drug, the solvent of a drug, or no drug. Samples were then cultured for ten more days for analysis.

### Mechanical characterization of agarose and tumor spheroids

To characterize the stiffness and stress-relaxation properties of hydrogels and tumor spheroids, milli-scale indentation and compression were performed, respectively, using a high-precision piezo-electric actuator-controlled system (CellScale, Canada). For indentation of hydrogel samples, indenters were constructed by gluing borosilicate glass spheres (radius R = 1.5 mm) to the ends of tungsten beams (Young’s modulus *E_b_* = 411 GPa) with circular crosssections. For parallel-plate compression of spheroid samples, stainless steel plates were fixed to the ends of the tungsten beams. The opposite end of the respective beam was then fixed onto a piezo-controlled positioning system.

A camera is positioned such that the displacement of the free end of the cantilevered beam, and hence displacement of the sample (indentation depth, d), was measured optically. Knowing the displacement of the piezo base (*z*) allowed the cantilevered beam deflection (*δ*) to be calculated as *δ* = *z* – *d*. Force (*F*) was determined from the deflection of the beam using the Euler-Bernoulli beam bending theory for a cantilevered beam loaded at the free end, which is dependent on other parameters of the beam including length (*L*), Young’s modulus (*E_b_*), and cross-sectional area moment of inertia (*I*), *F* = 3*δE_b_I/L*^3^. The force measurements were plotted against indentation depth. To estimate sample elastic indentation modulus (*E_i_*), the experimental force-indentation depth curves were fit to the Hertz contact equation, which estimates the force resulting from contact between a spherical body and a flat surface:

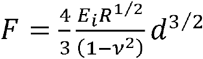

Here, *R* is the radius of the spherical indenter and ν is the sample’s Poisson’s ratio (approximated here to be 0.49, consistent with typical reported values for incompressible soft tissues [28–30]). The elastic indentation modulus of each sample was determined using a linear least squares method applied by a customized MATLAB code, which identified the value for modulus that minimized the error between the raw data and model Hertz curve.

For milli-indentation of agarose gels, indentation and retraction rates were kept constant at 4 μm/sec. To maintain elastic half-space assumptions, samples were indented to 10% of their measured height. All samples were glued to the stage to prevent floating and were tested in a PBS fluid bath at room temperature (~23 °C). At least three different locations were tested per sample.

For milli-compression of spheroids, samples were gently pipetted into the PBS fluid bath and tested at room temperature. A minimum of five spheroids were tested within each treatment group for each independent experiment. All spheroids were compressed to 20% deformation of their initial height, which induced a magnitude of force great enough to be captured by the cantilever beams according to their respective force resolutions while still maintaining elastic half-space assumptions [31]. A full test consisted of a compression phase, during which the deformation rate at the sample was 5 μm/s, followed by a 10-second hold phase in which the beam was held to a fixed sample deformation to assess the stress-relaxation response.

Soft tissues and their hydrogel simulants often exhibit viscoelastic responses to loads [28,32–34]. Multicellular tumor spheroids (MCTSs) in particular have been cited to viscously deform under small shear stresses in the span of several seconds [35,36]. As such, we held a constant deformation during mechanical testing in order to assess the stress-relaxation response (Supplementary Figure S2) and subsequently chose a set of parameters that could describe this viscoelastic behavior. The first of these properties, half-time relaxation (*t*_1/2_), defines the amount of time required to reach half of the total force that is relaxed within the sample, providing a measure of how quickly the sample relaxes. A second viscoelastic property we used was the ratio of steady state, relaxed modulus (*E_ss_*) to indentation modulus (*E_i_*). This property provided a measure for how much the sample relaxed, with lower ratios indicating a greater degree of relaxation. Steady state modulus (*E_ss_*) was found by substituting the steady-state, relaxed value of force obtained during the hold phase—at which point the sample had reached a new equilibrium—into the Hertz contact equation [28,29].

### Western blot

To assess the expression of the force generating protein myosin IIa, western blotting was performed. HCT116 and SUM149PT whole cell lysates were isolated from spheroids grown for seven days using mammalian protein extraction reagent (M-PER; ThermoFisher) following manufacturer protocols. Samples were treated with Bolt LDS Buffer (ThermoFisher) and Bolt reducing agent (ThermoFisher) and were heated to 70 °C for 10 minutes. Protein separation was done through electrophoresis using a Bolt 8% bis-tris gel (ThermoFisher) with wells loaded with 10 μg of protein. Subsequently, proteins were transferred onto a nitrocellulose membrane and then blocked overnight while shaken at 4 °C in TBST-5% BSA—a TBS Tween-20 (TBST; ThermoFisher) solution containing 5% bovine serum albumin (BSA; Sigma).

Primary antibodies for myosin IIa (1:5000; Cell Signaling Technology) and GAPDH (1:15000; Cell Signaling Technology) were diluted in TBST-5% BSA. The diluted primary antibodies were applied to appropriate sections of the membrane overnight while on a shaker at 4 °C. Membrane sections were washed overnight while shaken at 4 °C in TBST. Horseradish peroxidase-conjugated secondary antibodies (Cell Signaling Technology) diluted in TBST-5% BSA were then applied for 1 hour. Secondary antibody dilutions were as follows: myosin IIa at 1:10000 and GAPDH at 1:15000. Bound antibodies were detected using SuperSignal West Pico PLUS Chemiluminescent Substrate (ThermoFisher). Imaging was performed using a Chemidoc XRS+ system (BioRad). Quantification of band parameters were performed using the associated Image Lab software (BioRad).

### Contractility assay

To assess the contractile ability of the two cell lines, they were separately encapsulated in collagen gels and allowed to contract the gels over 24 hours. A collagen precursor solution was prepared from a monomeric collagen solution extracted from rat tails as described in Gong *et al*. [37]. The collagen precursor was neutralized following common protocols available for commercially available products with 10x PBS, 2 mM NaOH, and H_2_O while on ice. The neutralized collagen precursor was diluted with the appropriate media to a final concentration of 1.5 mg/mL. This solution was then used to resuspend a cell pellet of the respective cell line to achieve a cell density of 250,000 cells/mL. The cell suspension was deposited in a 24-well plate with 500 μL per well and allowed to gel for one hour in an incubator at 37 °C and 5% CO_2_. After gelation, 500 μL of media (with and without drugs or their solvents) was added to each well. Each gel was carefully detached from the walls of the well plate using a pipette tip and gentle shaking. The plate was cultured in the incubator for 24 hours, after which the cell-containing collagen gels were imaged. The areas of the gels were measured using ImageJ.

### Image analysis

CellProfiler was used to acquire the mean radius of samples grown without a vessel-like constraint [38]. A spherical shape was assumed, and the volume of the spheroids was calculated as 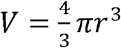.

ImageJ was used to acquire the major and minor axes of samples grown under a vessel-like constraint. For emboli constrained by the vessel-like constraint (with and without drugs), a cylindrical shape was assumed, and the volume of these emboli was calculated as *V* = *πr*^2^*l*, where *r* is the radius of the circular cross-section (minor axis) and *l* is the length of the major axis. For emboli that were not constrained by the vessel-like constraint, an oblate ellipsoidal shape was assumed, and the volume of the emboli was calculated as 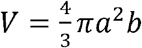, where *a* is the radius of the major axis and *b* is the radius of the minor axis.

### Statistics

All scatter plots and bar graphs are reported as mean ± standard error of the mean and were plotted using OriginPro (OriginLab). Statistical differences between the growth conditions via constraint or drug inhibition were determined by one-way ANOVA tests with Tukey post hoc using OriginPro (OriginLab). Differences were considered significant at p < 0.05.

## RESULTS

### Fabrication of microchannels as a vessel model in stiff or soft hydrogels

Vasculature can be generalized as tubular organs within the connective tissue of the body; thereby, we modeled vessels as cylindrical channels within a hydrogel matrix. To construct the vessel model, a microchannel was formed by templating a rod within an agarose matrix. The rod had a diameter of 200 μm, which is approximately the diffusion limitation of oxygen in tissue [39,40] and is similar in diameter to that of collecting lymphatic vessels [41,42]. Two variants of a 3D-printed frame were used to suspend the rod: a short template for the *soft* agarose and a long template for the *stiff* agarose (Figure 1A and Supplementary Figure S1, respectively). The integrity of the microchannel was confirmed by imaging of a test sample embedded with fluorescent nanoparticles (Figure 1B). The hydrogel matrix used was agarose, which was chosen for its bioinert properties that prevent cell adhesion and degradation, its nanoporosity that prevents cell migration and invasion, and its wide range of stiffness that may be tuned by changing the agarose concentration [18,19,43]. Agarose concentrations of 1.0% (w/v) and 0.3% (w/v) were used to mimic excessive (*stiff*) and normal (*soft*) tissue stiffness, respectively (Figure 1C). The elastic modulus of agarose at 1.0% was measured to be 6130 ± 213 Pa, while 0.3% agarose had an elastic modulus of 302 ± 11 Pa. We used the vessel model developed here to isolate the effects of the geometric constraint and mechanical stiffness on the growth of tumor emboli without having to account for matrix modifications beyond those due to tumor volumetric growth.

### Stiffness of the vessel-like constraint influences tumor emboli morphology

Using both human colon cancer cells (HCT116) and human breast cancer cells (SUM149PT), we first compared the growth of tumor spheroids (no mechanical constraint) to the growth of tumor emboli (in the vessel-like constraint model) in either *stiff* or *soft* agarose (Figure 1C). We first noted that both cell lines had similar doubling times (Supplementary Figure S3). Growing as spheroids, both cell lines maintained a projected circular shape, commonly seen in other studies (Figures 2A and 2B) [17,44–46]. Both cell lines also produced spheroids that exhibited similar growth dynamics, following the characteristic Gompertzian, or sigmoidal, volumetric growth rate model [19]. The Gompertzian growth curve is characterized by an exponential growth period followed by a plateau, which was reached here after approximately two weeks by both cell lines at tumor spheroid diameters of 719 ± 53 μm and 559 ± 33 μm, for the HCT116 and SUM149PT cell lines, respectively. The plateau in growth has been attributed to the inability for nutrients to diffuse into and waste to diffuse out of larger spheroids, which limits their growth [47,48].

**Figure 2.**
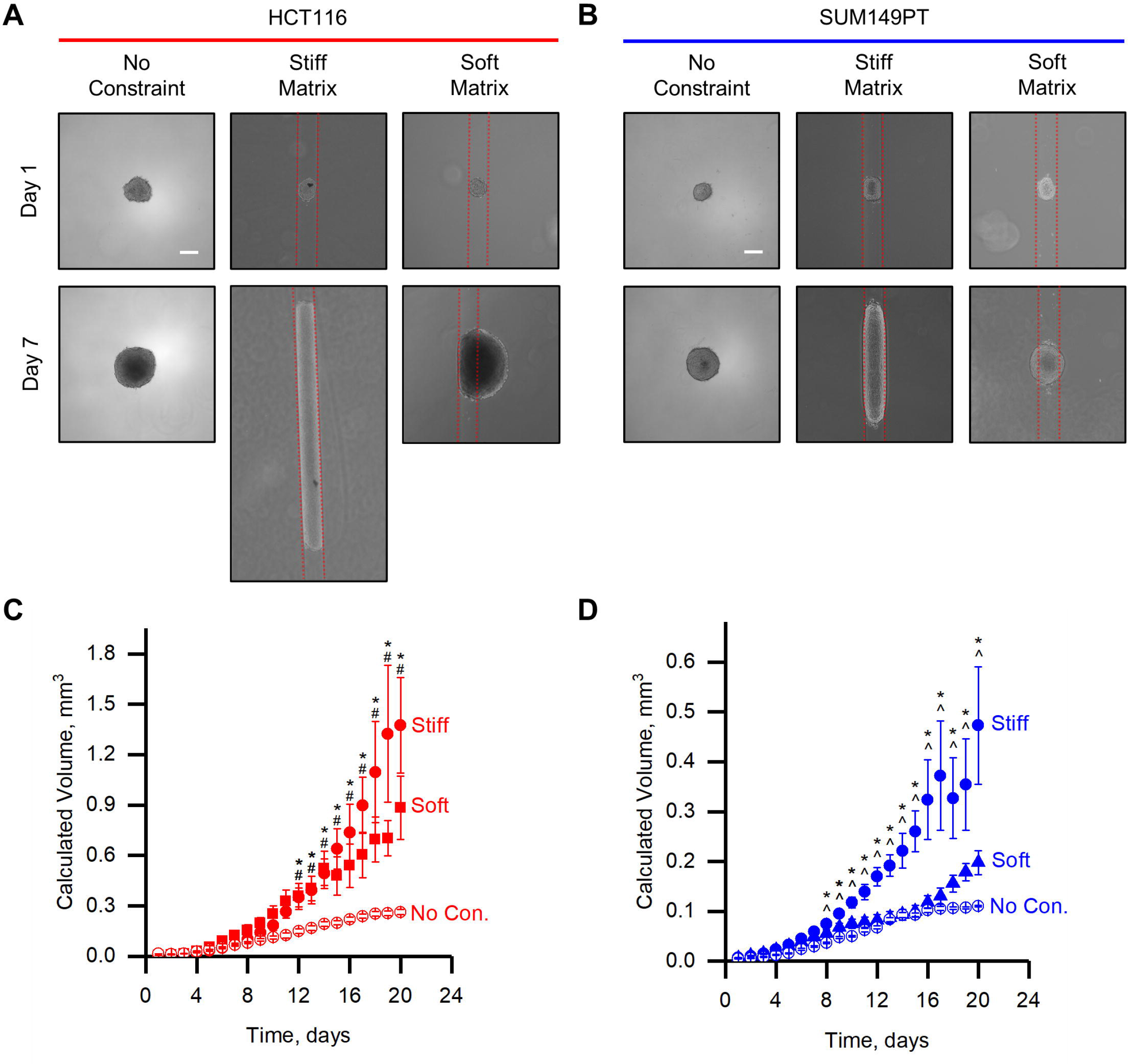
Stiffness of the vessel-like constraint influences tumor emboli growth and morphology. A) Phase contrast images of HCT116 cells grown without constraint as spheroids (left), within the stiff vessel model (middle), and within the soft vessel model (right). B) The same as A) except for SUM149PT cells. C) Calculated volumes over a growth period of 20 days of HCT116 spheroids (No Con.) and emboli in both the soft matrix (Soft) and stiff matrix (Stiff). D) The same as C) but for SUM149PT cells. Scale bars in A) and B) are 200 μm. All data points are presented as mean ± SEM. * denotes p < 0.05 for Stiff compared to No Con. # denotes p < 0.05 for Soft compared to No Con. ^ denotes p < 0.05 for Stiff compared to Soft.

When grown as emboli within the *stiff* vessel-like constraint model (stiff vessel model), the emboli were confined to and elongated within the tubular geometry of the microchannel (Figure 2A and 2B, Supplementary Figure S4B). The stiff matrix restricted the expansion of the HCT116 tumor emboli circumferentially, forcing the cells to grow along the channel length. The SUM149PT tumor emboli were also constrained circumferentially, although they bulged slightly against the stiff matrix. The elongated tumor emboli in the stiff vessel model often exceeded the length of the short template, necessitating the longer template to be able to measure growth over the entire culture period. The volumetric growth curves for either cell line in the stiff vessel model do not reach an obvious plateau after 20 days of growth, and the emboli volumes significantly exceeded those of the spheroids grown without constraint (Figure 2C and 2D). Furthermore, the HCT116 emboli volumes significantly exceeded those of the SUM149PT emboli. This geometry seems to have conferred a growth advantage to the cells, likely because the 200 μm-diameter of the channel physically restricted the tumor emboli from growing larger (in the cross-section of the channel) than the diffusion limit distance.

The modulus of most healthy soft tissue is below 1 kPa: normal colon tissue has been found to possess a steady state modulus of ~700 Pa and normal breast tissue possess a Young’s modulus of also ~700 Pa under low stress tests [49,50]. As the elastic modulus of the *stiff* agarose exceeds that of most healthy soft tissues, we repeated the experiment but with cylindrical channels molded into *soft* agarose that are closer to normal tissue stiffness [51]. Interestingly, in the *soft* vessel-like constraint model (soft vessel model), emboli growth was not confined to the cylindrical channel as in the stiff vessel model. Rather, after forming a spheroid that filled the diameter of the channel, the tumor emboli grown from both cell lines eventually bulged into the matrix (Figures 2A and 2B). Although they were growing against the soft matrix and did not significantly elongate along the channel, the emboli in the soft vessel model from both cell lines had intermediate growth rates between the unconstrained spheroids and emboli in the stiff vessel model (Figures 2C and 2D). They also did not reach an obvious plateau in their growth after 20 days.

The growth dynamics and tumor emboli shapes, similar to the stiff vessel model, were cell-line dependent. HCT116 emboli grown in the soft vessel model increased in volume faster than when grown as spheroids, but slower than when grown as emboli in the stiff vessel model (Figure 2C). In contrast, SUM149PT emboli grown in the soft vessel model did not increase in volume significantly faster than when grown as spheroids until the time that the spheroids reached their plateau size (Figure 2D). The SUM149PT emboli grown in the stiff vessel model, however, still increased in volume fastest as compared to the emboli in the soft vessel model and spheroids. The shape to which the emboli grew were also different for the two cell lines. The HCT116 emboli expanded from the channel into a roughly oblate ellipsoidal shape (Figure 2A, projection, and Supplementary Figure S4A); whereas the SUM149PT emboli maintained a more circular projection throughout the growth period (Figure 2B). Together, we demonstrated that imposing a vessel-like geometrical constraint on the growth of tumor cell aggregates significantly impacted the rates and shapes of their growth. In addition to the geometrical constraint, the stiffness of the constraint was a determinant of the rate and shape of tumor cell aggregate growth, demonstrating an apparent mechanosensitivity of the tumor cells.

As the overall shapes of the tumor emboli grown within the soft vessel model were drastically different, we characterized their morphologies as a function of time. Examples of projected tumor outlines are plotted as heat maps as a function of time in the insets of Figures 3A and 3B for HCT116 and SUM149PT cells, respectively. After filling the microchannel, the HCT116 emboli elongated to a relatively high aspect ratio of between 2 to 5 along the length of the soft vessel model. The elongation continued until around day 7, when they bulged out of the channel and into the agarose gel, reducing their aspect ratio (Figure 3A). In contrast, the SUM149PT cells maintained a uniform, round projection seemingly without being influenced by the vessel-like geometry as their aspect ratio was almost 1 throughout the culture period (Figure 3B). These observations suggest that the HCT116 emboli were more sensitive to the geometry of the soft vessel model—initially growing along the open channel—as compared to the SUM149PT emboli that appeared unaffected by the soft constraint.

**Figure 3.**
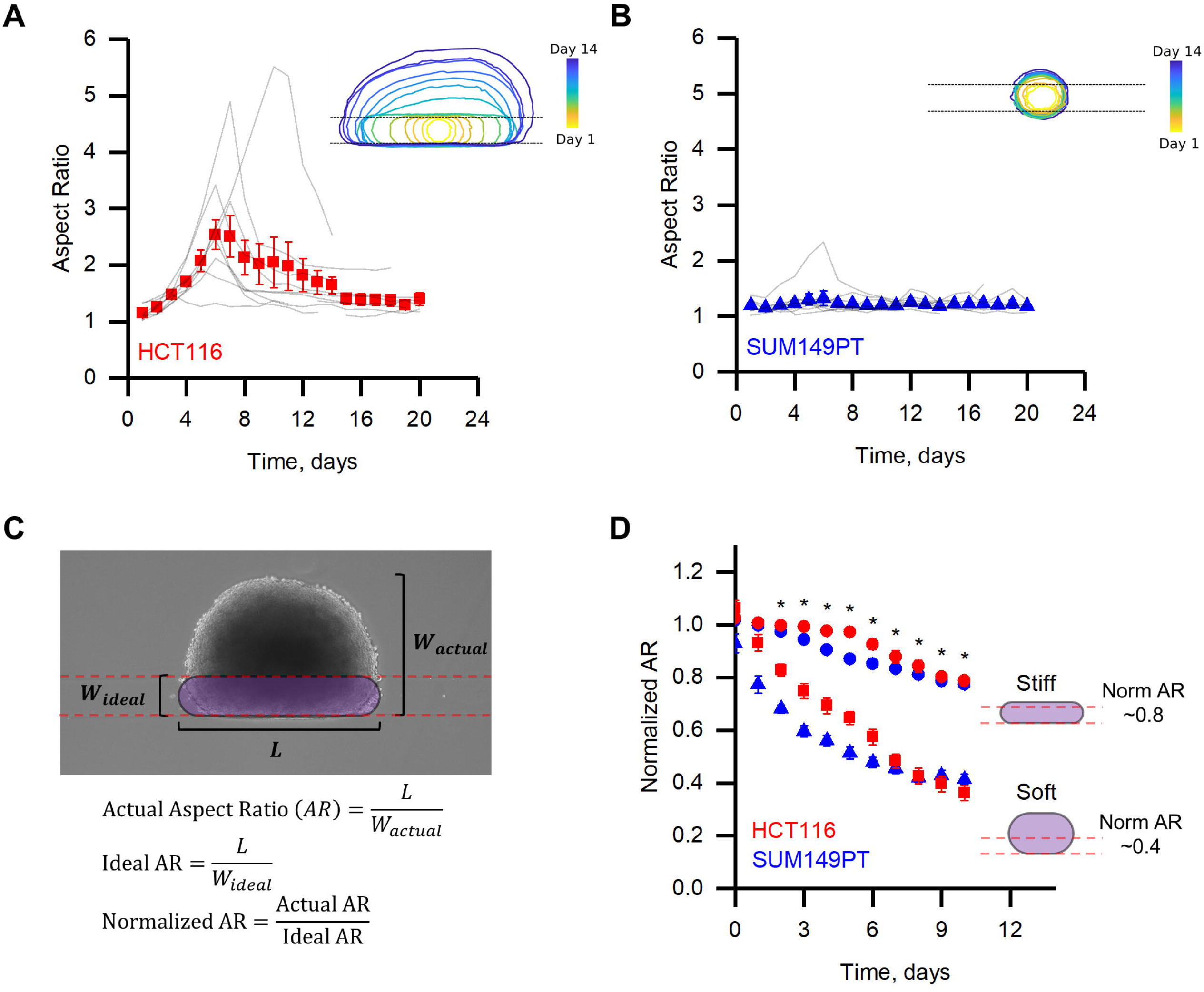
Mechanosensitivity to a soft vessel-like constraint leads to differences in tumor emboli growth and morphology. Average aspect ratio of A) HCT116 tumor emboli (red, squares) and B) SUM149PT tumor emboli (blue, triangles) grown in soft vessel models overlaid onto aspect ratios of individual tumor emboli (grey) over 20 days. Inset diagrams depict the morphology of average tumor emboli, of the respective cell type, over a 14-day period. C) Schematic illustrating the “normalized” aspect ratio. The ideal width (*w_ideal_*) is set to be the diameter of the vessel-like constraint (200 μm) and the length (*L*) is the major axis. D) Normalized aspect ratios of both HCT116 and SUM149PT emboli grown in soft and stiff vessel models (HCT116, red: stiff = circles, soft = squares; SUM149PT, blue: stiff = circles, soft = triangles). At least five samples (n ≥ 5) are depicted for all time points in each graph. All data points are presented as mean ± SEM.

To quantify the amount of bulging into the matrix, we defined a normalized aspect ratio to compare the actual bulged shape to an ideal, channel-confined tumor (Figure 3C). The ideal aspect ratio is the length that the tumor grew along the channel divided by the channel width. The actual aspect ratio is the length that the tumor grew along the channel divided by its actual width. The normalized aspect ratio (actual/ideal) decreases with increased bulging against the microchannel. Plotting the average normalized aspect ratio for emboli from each cell line in soft and stiff vessel models (Figure 3D) highlights our observation that in no case was bulging fully restricted. The stiffer matrix, however, provided significantly more constraint. Also clearly illustrated is the immediate bulging of the SUM149PT emboli grown in the soft vessel model (Figure 3D). This analysis of the time-course of the shape development of the tumor emboli in the soft vessel-like constraint model reveals that HCT116 emboli were initially sensitive to the mechanical constraint, while the SUM149PT emboli were not.

### Tumor spheroid biomechanical properties are products of force-generating mechanisms

To account for the dissimilarity in the evolution of tumor emboli shapes between cell lines, we hypothesized that their elastic moduli were significantly different and, therefore, they possessed different magnitudes of mechanical mismatch with the matrix of the vessel models (*E_ves_/E_emb_* ≠ 1). In fully embedded tumor models, this mechanical mismatch has been shown to drive the shape of tumor cell aggregate growth [18]. Specifically, when the embedded tumor is softer than the hydrogel matrix, it will grow into an ellipsoid; whereas, when the embedded tumor is stiffer than the hydrogel matrix, it will grow into a sphere. Therefore, we hypothesized that the SUM149PT spheroids were stiffer (0 < *E_ves_/E_emb_* < 1), whereas the HCT116 spheroids were softer (*E_ves_/E_emb_* > 1), than the soft agarose.

We measured the mechanical properties of HCT116 and SUM149PT tumor spheroids after 10 days of growth using parallel-plate compression. Indeed, the measured elastic modulus of the HCT116 (*E_emb, HCT_* = 117 ± 31 Pa) and SUM149PT (*E_emb, SUM_* =1195 ± 332 Pa) spheroids (Figure 4A) seemingly supported our hypothesis. In the stiff vessel model, the mechanical mismatch between the vessel matrix and the emboli would be *E_ves_/E_emb_* = 52 for the HCT116 and 5.1 for the SUM149PT emboli. In the soft vessel model, the HCT116 emboli were initially confined to the soft vessel, *E_ves_/E_emb_* = 2.6, while the SUM149PT were not, *E_ves_/E_emb_* = 0.25. After an average of about seven days of growth in the soft vessel model, when they start to bulge into the matrix, it is likely that the HCT116 tumor emboli stiffened in response to the constraint. We and others have observed increased expression of actin surrounding tumor cell aggregates when they are embedded in hydrogels with elastic moduli similar to or stiffer than the aggregates, but not when the hydrogels are softer (Supplementary Figure S5, Taubenberger *et al*., 2019). The development of such a thick actin shell has been shown to lead to an increase in aggregate stiffness [52,53].

**Figure 4.**
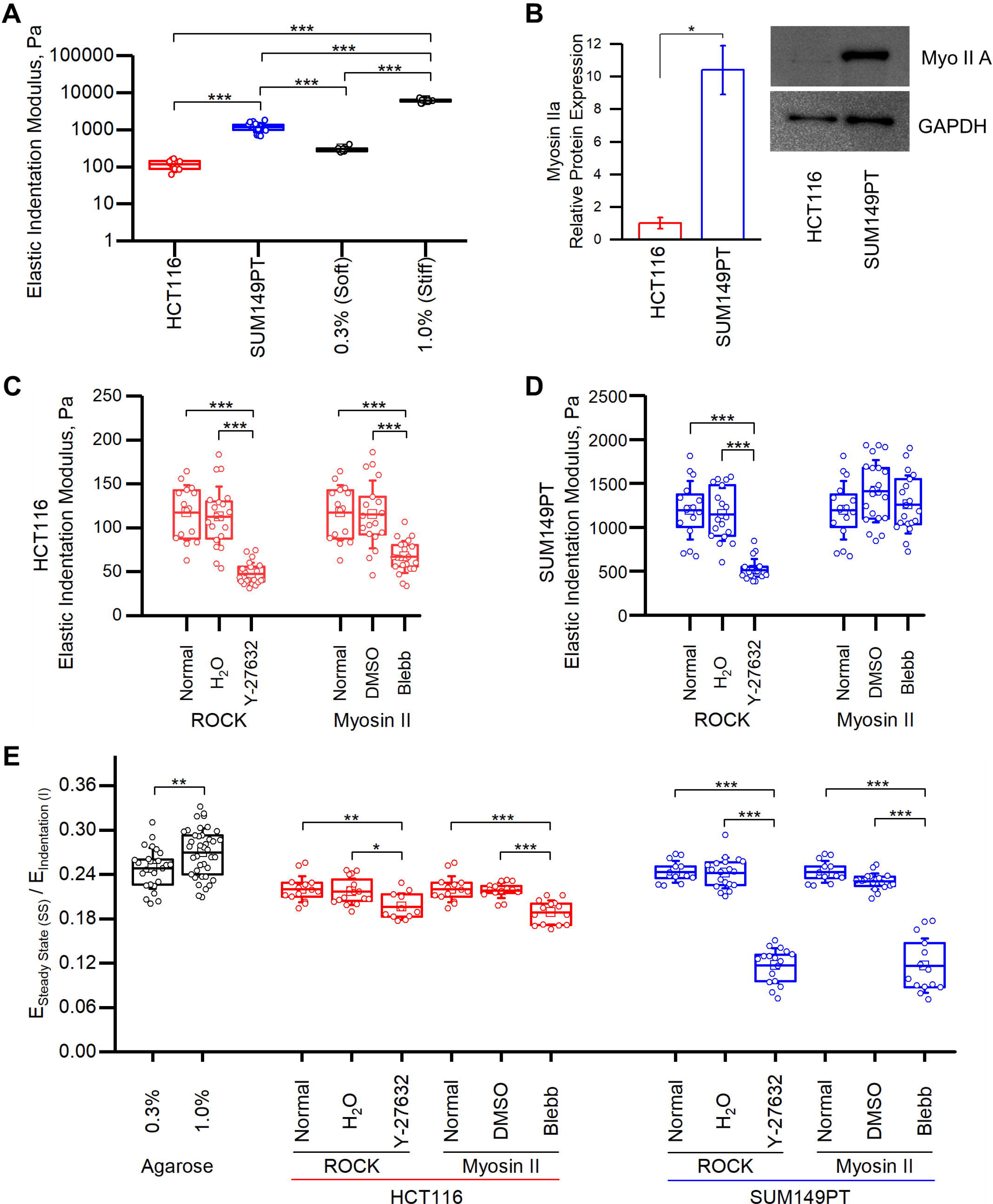
Inhibition of force-generation leads to shifts in tumor spheroid biomechanics. A) Elastic moduli of HCT116 and SUM149PT spheroids grown for seven days compared to the stiffness of the *soft* and *stiff* vessel-like constraint models. B) Relative expression and representative images of western blots of myosin IIa and GAPDH from HCT116 and SUM149PT spheroids over three independent experiments. C) Elastic indentation moduli of HCT116 spheroids grown with and without drugs after seven days of treatment. D) Same as (C), but for SUM149PT. E) Ratio of steady state moduli over elastic indentation moduli of agarose as compared to HCT116 and SUM149PT spheroids with and without drug treatments. Box plots are representative of three independent experiments with n ≥ 5 samples each. *, **, and *** denotes p < 0.05, p < 0.01, and p < 0.001, respectively.

Cell aggregate stiffness has been linked to (i) actomyosin-generated tension in embryonic tissues and in epithelial sheets as well as to (ii) excessive extracellular matrix deposition and crosslinking in disease processes such as cancer [54–58]. Cells in spheroid culture produce ECM proteins [36,59,60]; however, they are not thought to be rigid and load-bearing [36]. Therefore, we concluded that tension generation via actomyosin contractility was likely the major determinant of the stiffness of the tumor spheroids. Seeming to corroborate this conclusion, western blot analysis showed that the myosin IIa expression of SUM149PT cells was ten times higher than that of HCT116 cells (Figure 4B) and SUM149PT cells exhibited significantly higher force generation in a collagen-gel contraction experiment (Supplementary Figure S6A).

To explore whether actomyosin contractility significantly contributes to the stiffness of the tumor spheroids, we tested whether their stiffness could be modulated via perturbations of myosin-related force generation through two commonly targeted routes: the Rho-associated protein kinase (ROCK) pathway and myosin II. The ROCK pathway is associated with regulation of cell shape, movement, and contractility by phosphorylation of myosin light chains, which increases myosin II ATP activity [61–63]. Myosins are motor proteins primarily associated with contraction and motility when they work in concert with actin filaments. Myosin II is specifically responsible for cellular contraction [64–66]. Spheroids that had stably aggregated after three days were treated with drugs targeting either ROCK—using Y-27632 (100 μM)—or myosin II directly—using blebbistatin (10 μM). The growth of the tumor spheroids was monitored over seven days after treatment and then their stiffness was measured. The growth of the HCT116 spheroids was not affected by either inhibitor as all spheroids reached diameters on the same order as the normal, untreated spheroids (Supplementary Figure S7A). In terms of their stiffness, however, both the ROCK inhibitor (Y-27632) and blebbistatin significantly reduced their average stiffness by 60% and 43%, respectively (Figure 4C). In contrast, both inhibitors hindered the growth of the SUM149PT spheroids, as these samples reached diameters about 100-μm smaller than untreated and control samples (Supplementary Figure S7B). The introduction of Y-27632 reduced the average stiffness of the SUM149PT spheroids by 57%, similar to the effects on HCT116 spheroids. However, blebbistatin interestingly had no effect on the average stiffness of the SUM149PT spheroids (Figure 4D). A similar observation of the effects of blebbistatin on SUM149PT cells was made in a collagen-gel contraction experiment: it did not inhibit contractility of SUM149PT cells whereas the ROCK inhibitor did (Supplementary Figure S6B). Together, these results indicate that inhibition of myosin-related force generation through either the ROCK pathway or myosin II decreases the elastic moduli of tumor spheroids, with the exception of blebbistatin-treated SUM149PT spheroids, which were not significantly affected.

### Myosin inhibition increases stress relaxation in tumor spheroids

In addition to effects on the elastic moduli of the spheroids, treatment of the spheroids with the ROCK inhibitor and blebbistatin also altered their viscoelastic properties. We used the stress relaxation behavior of the agarose and MCTSs to quantify their steady-state, relaxed modulus, and half-time of relaxation. The steady-state elastic modulus of both the agarose hydrogels and the untreated MCTSs was between 22% and 27% of the indentation modulus (*E_ss_/E_i_*, Figure 4E). For HCT116 spheroids treated with either inhibitor, the steady-state modulus was slightly, although significantly, decreased with respect to the indentation modulus. In contrast, the SUM149PT spheroids treated with either inhibitor displayed a large drop in steadystate modulus compared to the untreated spheroids. The ratio *E_ss_*/*E_i_* for the SUM149PT spheroids was decreased to 0.12 ± 0.04 and 0.12 ± 0.02 for the blebbistatin and ROCK inhibitor, respectively, from 0.24 ± 0.01 for the untreated SUM149PT spheroids. Taking into account the relaxation in the mechanical mismatch for one of these cases, SUM149PT spheroids treated with the ROCK inhibitor, we observed that this increased relaxation switched *E_ves_/E_emb_* for the soft vessel model from less than one, (*E_ves_/E_emb_*)*_i_* = 0.59, to greater than one, (*E_ves_/E_emb_*)*_ss_* = 1.2. The relaxation rates of all MCTSs were also increased with either drug treatment, but the relaxation half-times remained within the range of 1.7 to 2.0 seconds for all MCTSs (both untreated and drug treated) whereas for agarose relaxation half-times were between 30 and 50 seconds (Supplementary Figure S8). Thus, in addition to decreasing the elastic moduli of most tumor spheroids, the ROCK and myosin II inhibitors increased the magnitude and rate of stress relaxation of all tumor spheroids. The one case where no effect was made on elastic modulus was SUM149PT spheroids treated with blebbistatin.

### Inhibition of force generation leads to constraint of emboli growth by the soft vessel model

Since the ROCK and myosin II inhibitors increased the compliance and/or the degree of relaxation of the HCT116 and SUM149PT spheroids, they effectively increased the magnitude of mechanical mismatch between the spheroids and the matrix of the soft vessel-like constraint model. We therefore hypothesized that treatment of the emboli with these inhibitors would lead to increased confinement of growth along the *soft* vessel model. To test this hypothesis, the inhibitors were introduced to tumor emboli within the soft vessel model once they grew to the diameter of the microchannel. Emboli growth and morphologies were measured and monitored over a period of ten days.

When treated with either inhibitor, tumor emboli from both cell lines displayed increased confinement, to different degrees, of growth along the soft vessel model. Treatment of HCT116 emboli with the ROCK inhibitor lessened emboli bulging in the soft vessel model as compared to untreated (normal) emboli, with normalized aspect ratios of 0.70 ± 0.02 and 0.48 ± 0.08 on Day 7, respectively (Figure 5B). In contrast, blebbistatin treatment recovered the level of HCT116 emboli confinement to that of the untreated stiff vessel model (Figure 5C). The drug treatment did not appear to affect the growth of the tumor emboli as their calculated volumes were not significantly affected (Supplementary Figures S7C and S7E).

**Figure 5:**
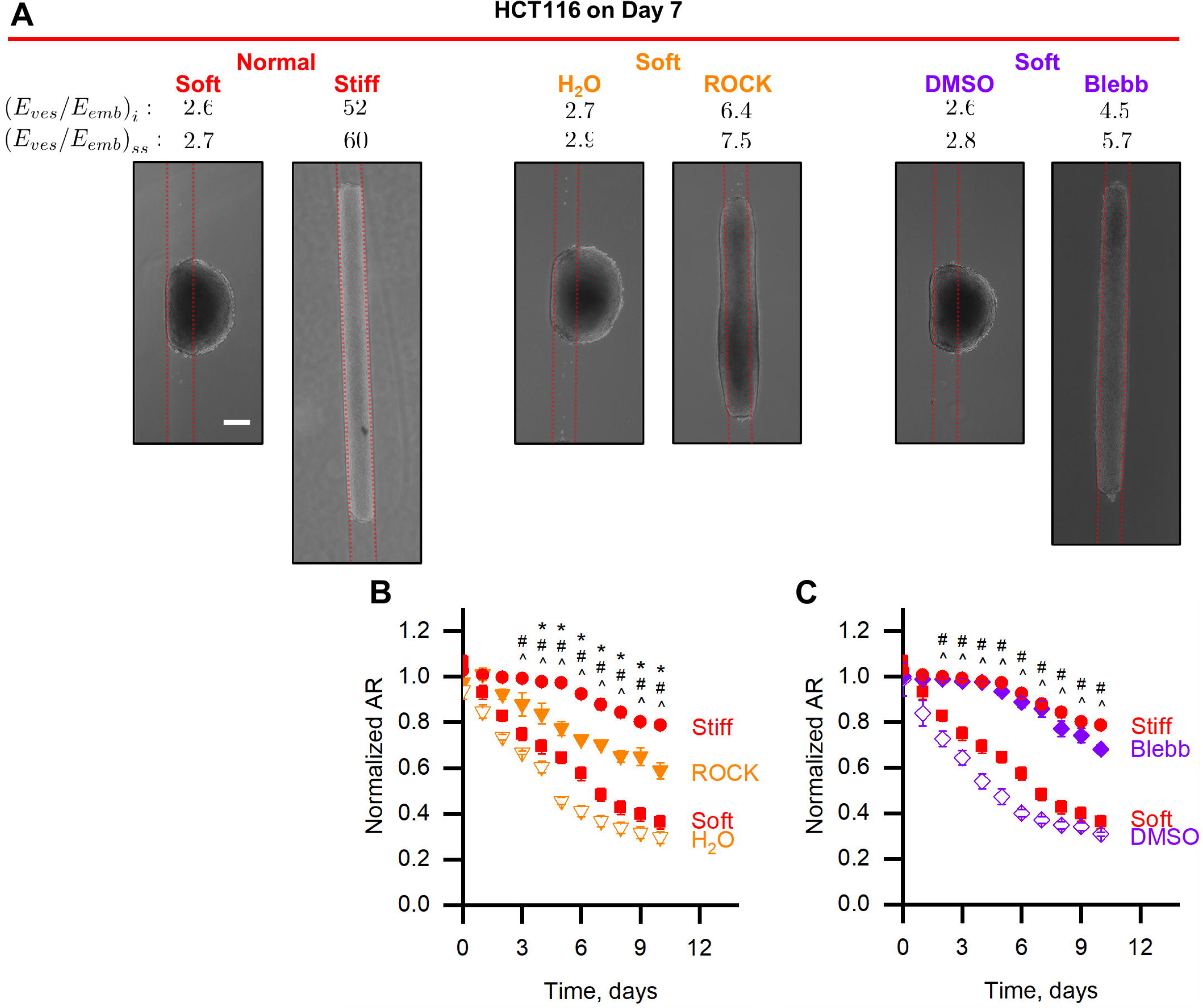
ROCK and myosin II inhibition lead to increased confinement of HCT116 emboli in the soft vessel model. A) Representative phase contrast images of HCT116 emboli grown without drug treatment (Normal) or with a drug (and compared with its control). All images taken on Day 7. Red dotted lines denote the boundaries of the vessel-like constraint. Scale bar is 200 μm. B) Normalized aspect ratio of HCT116 emboli treated with the ROCK inhibitor (orange triangles) compared to those grown under soft and stiff vessel-like constraints (red squares and circles, respectively). C) Same as B), but with blebbistatin (purple diamonds). Data is representative of n ≥ 3. *denotes p < 0.05 for drug-treatment compared to Stiff. ^ denotes p < 0.05 for drug-treatment compared to Soft. # denotes p < 0.05 for drug-treatment compared to control.

Growth of the SUM149PT emboli in the soft vessel model was more sensitive to the ROCK inhibitor than blebbistatin (Figure 6A). ROCK inhibitor-treated SUM149PT emboli grew along the soft vessel model similarly to untreated emboli in the stiff vessel model (Figure 6B). On the other hand, the SUM149PT emboli treated with blebbistatin appeared to be significantly less affected by the drug, forming emboli with normalized aspect ratios similar to those of the untreated and control samples in the soft vessel model (Figure 6C). However, a slight elongation along the microchannel was observed under the treatment with blebbistatin (Figure 6A). As with the HCT116, the calculated volumes of the SUM149PT emboli were not significantly affected (Supplementary Figures S7D and S7F). Together, we demonstrated that the increase in the mechanical mismatch between the soft vessel model matrix and the tumor emboli—due to decreased stiffness *or* increased relaxation of the tumor cell aggregates after treatment with a ROCK or myosin II inhibitor—increased the confinement of emboli growth in the soft vessel model. In two cases (ROCK-treated SUM149PT and blebbistatin-treated HCT116) the level of confinement was nearly as high as that provided by the stiff vessel model.

**Figure 6:**
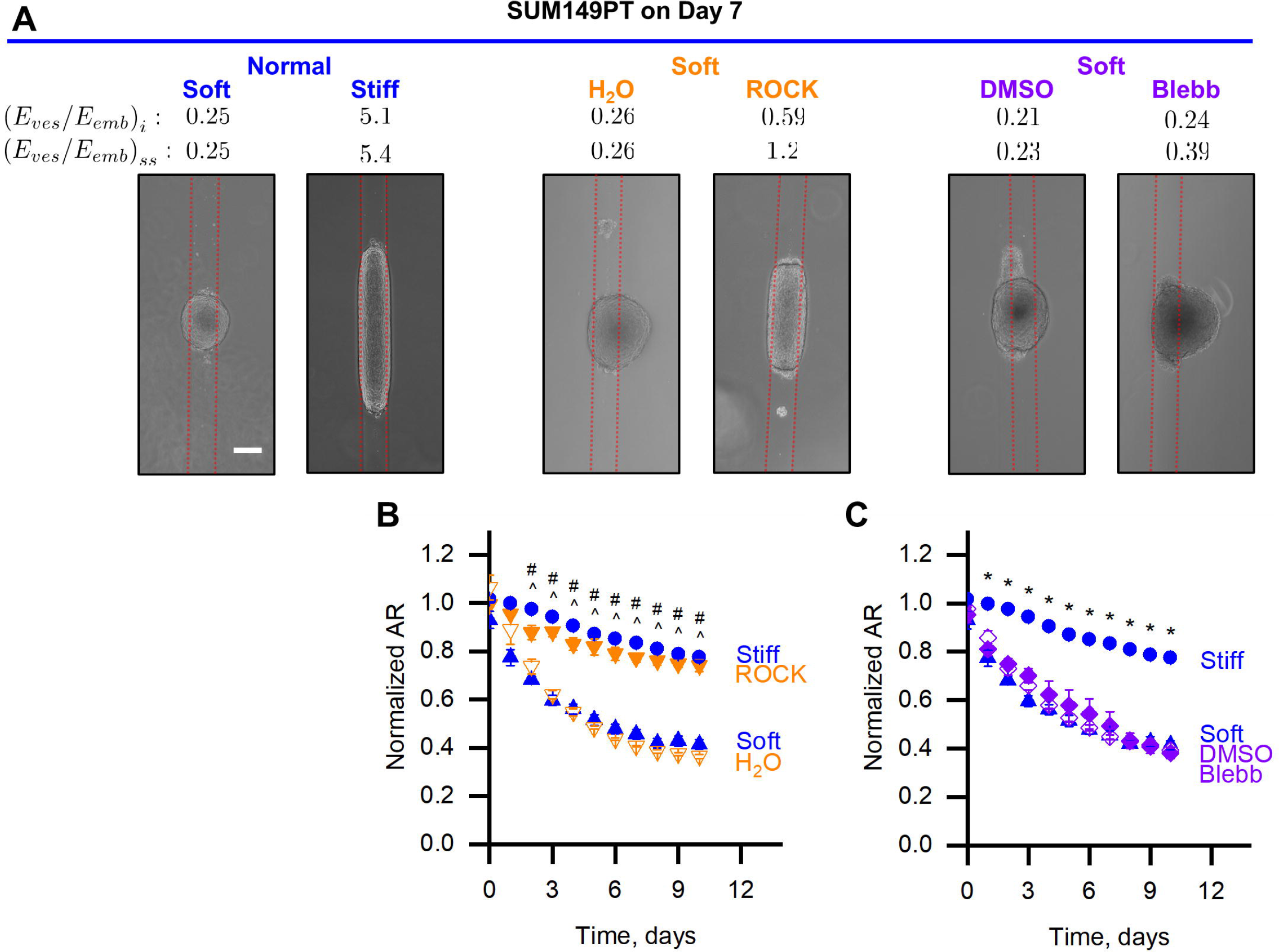
ROCK inhibition, but not myosin II inhibition, drives the confinement of SUM149PT emboli growth in the soft vessel model. A) Representative phase contrast images of SUM149PT emboli grown without drug treatment (Normal) or with a drug (and compared with its control). All images taken on Day 7. Red dotted lines denote the boundaries of the vessel-like constraint. Scale bar is 200 μm. B) Normalized aspect ratio of SUM149PT emboli treated with the ROCK inhibitor (orange triangles) compared to those grown under soft and stiff vessel-like constraints (blue triangles and circles, respectively). C) Same as B), but with blebbistatin (purple diamonds). Data is representative of n ≥ 3. * denotes p < 0.05 for drugtreatment compared to Stiff. ^ denotes p < 0.05 for drug-treatment compared to Soft.

## DISCUSSION

In this study we examined the morphological response of a tumor cell aggregate growing within a vessel-like constraint as it would in the context of a tumor embolus. We demonstrated how the magnitude of the mechanical mismatch between the vessel-like constraint and the tumor emboli (*E_ves_/E_emb_* ≠ 1) can lead to drastically different tumor shapes. Here, the mechanical properties of the vessel-like constraint (the agarose matrix) in relation to that of the tumor emboli directed tumor emboli to either elongate along the microchannel length (*E_ves_/E_emb_* > 1) or bulge against the constraint (0 < *E_ves_/E_emb_* < 1). The introduction of inhibitors that target myosin-related force generation decreased the elastic modulus and/or increased the stress relaxation of the tumor cell aggregates, effectively increasing the mechanical mismatch between the vessel-like constraint and the emboli. The increased mechanical mismatch after drug treatment was correlated with increased confinement of tumor emboli growth in the soft vessel model.

The confinement of tumor emboli to grow within a vessel geometry—either due to an excessively stiffened matrix or weak actomyosin contractility of the tumor cells—may provide a growth advantage to the emboli. Indeed, when confined within the *stiff* vessel model, emboli from both cell lines (*E_ves_/E_emb_* = 52 for the HCT116 and 5.1 for the SUM149PT) grew dramatically larger in volume with no plateau in growth, in contrast to their unconstrainted MCTS counterparts. The reasons for the dramatic growth were likely access to nutrients and space for uninhibited growth. The confinement of radial expansion imposed by the stiff matrix on emboli from both cell lines ensured that no point within the cylindrical tumor (200 μm diameter) was more than 100 μm from the tumor-matrix boundary, well within the diffusion limit (~200 μm) of oxygen in tissues [67]. In addition to proximity to oxygen and nutrients, the cells had an open channel into which they could expand, uninhibited.

Strikingly, even with an open channel into which they could expand, tumor emboli grown within the *soft* vessel model bulged, eventually, into the matrix. Emboli morphology was determined by the mechanical mismatch between the matrix of the soft vessel model and the emboli. HCT116 aggregates that were slightly more compliant than the matrix of the soft vessel model *E_ves_/E_emb_* = 2.6 were initially confined to the soft vessel model, but after 7 days bulged into the matrix and formed oblate ellipsoids. On the other hand, SUM149PT aggregates that were much stiffer than the matrix of the soft vessel model *E_ves_/E_emb_* = 0.25 maintained a spherical shape, apparently unaffected by the vessel-like constraint. This observed phenomenon has been predicted by calculations of the elastic strain energy stored in a system composed of an infinite isotropic matrix containing an inhomogeneous ellipsoidal inclusion [68]. If the inclusion is subjected to an isotropic growth strain, the elastic strain energy is minimized by a flat ellipsoidal inclusion when the inclusion is more compliant than the matrix (*E_mat_/E_inc_* > 1), and a sphere will minimize the elastic strain energy when the inclusion is stiffer than the matrix (0< *E_mat_/E_inc_* < 1). In a previous study we showed that tumor cell aggregates grown in agarose hydrogels display the same strain-energy minimizing growth behavior as predicted by the aforementioned calculations [18].

The delayed bulging of the HCT116 tumors into the matrix of the soft vessel model may be due to the development of a shell of F-actin surrounding the tumor aggregates, the thickness of which increases with the stiffness of the matrix in which the aggregate is embedded (Supplementary Figure S5) [52]. The development of the F-actin shell, suggesting a mechanosensitive response of the aggregate to confinement, has been shown to correlate with an increase the stiffness of the aggregate [52,69]. This switch from confinement to bulging may only have been observed because the HCT116 spheroids were so close in elastic modulus (117 ± 31 Pa) to the soft agarose (302 ± 11 Pa), such that the level of increase in stiffness accompanying the development of a thick F-actin shell could have inverted the mechanical mismatch. Similarly, the inability of the stiff vessel model to fully constrain the tumor emboli at later time points may be attributable to the development of a thick F-actin shell and the associated actomyosin contractility.

We found that the difference in mechanical properties between the spheroids derived from the two cell lines was correlated with differential expression of myosin IIa, as seen in the western blot (Figure 4B), and accompanying increased actomyosin contractility, as see in the contractility assays (Supplementary Figure S6). Myosin II plays a major role in the contractility of the cytoskeleton, enabling cells to generate forces for migration, remodeling, and the innate stiffness of cells and tissues [58,70–72]. Inhibition of myosin—either via ROCK (Y-27632), for which myosin II is a downstream effector, or directly (blebbistatin)—decreased the stiffness and/or increased the relaxation of tumor spheroids. The ROCK inhibitor decreased the elastic modulus of spheroids from both cell lines, which has been observed for other cells and tissues [52,55,73]. Blebbistatin, however, only decreased the elastic modulus of the HCT116 spheroids, not the SUM149PT spheroids. The SUM149PT cells expressed myosin II at a much higher level, which perhaps we were not able to effectively inhibit using the limited blebbistatin concentration attainable due to its poor solubility in water [66]. Notably, although the elastic modulus of the SUM149PT spheroids was not affected by blebbistatin, its relaxed, steady-state modulus was drastically affected. A large drop in the relaxed, steady-state modulus was also observed for the SUM149PT spheroids treated with the ROCK inhibitor.

Introducing the inhibitors of force-generation into emboli cultures produced drastic changes in the morphology of the tumor emboli. Both inhibitors drove the HCT116 growth along the *soft* vessel-like channel. However, Y-27632 was less efficacious as the emboli still slightly bulged into the matrix and the normalized aspect ratio was not as large as that for emboli treated with blebbistatin, which matched that of untreated HCT116 emboli grown within the *stiff* vessel model. Both of the inhibitor treatments reduced the elastic moduli of HCT116 spheroids to well below the soft agarose elastic (or steady-state) modulus (*E_ves_/E_emb_* = 6.4 for the ROCK inhibitor and 4.5 for blebbistatin). With such a significantly increased modulus mismatch between the soft vessel matrix and the ROCK-treated HCT116 spheroids, it is not clear why the emboli still bulged into the matrix. One possibility may be the activation of complementary pathways to ROCK, such as Rac1 and CDC42, that have the same downstream effectors, such as myosin II, enabling the HCT116 emboli to elicit force-generating activities [62,74]. Blebbistatin, on the other hand, blocks myosin II activity regardless of the upstream activator.

The SUM149PT tumor emboli also exhibited interesting morphological responses to the drug treatments: Y-27632 was most efficacious at driving emboli growth along the soft vessel-like channel, while blebbistatin appeared to have a minimal effect. Y-27632 reduced the elastic modulus of SUM149PT spheroids, but it was still significantly greater than the elastic modulus of the soft agarose ((*E_ves_/E_emb_*)*_i_* = 0.59, compared to 0.25 for untreated). Interestingly, because the drugs had the effect of drastically increasing the relaxation of the SUM149PT spheroids, the relaxed, steady-state modulus of the SUM149PT spheroids was just less than the steady-state modulus of the soft agarose, inverting the mechanical mismatch ((*E_ves_/E_emb_*)*_ss_* = 1.2). On the other hand, blebbistatin did not affect the elastic modulus of the SUM149PT spheroids, and even taking into account the relaxation, the relaxed modulus was still significantly greater than the soft agarose ((*E_ves_/E_emb_*)*_ss_* = 0.39). However, this reduction in steady-state modulus did appear to allow the SUM149PT emboli in the soft vessel model to initially elongate slightly along the length of the channel, which the untreated emboli did not do.

In summary, we found that the SUM149PT spheroids were approximately 10 times stiffer than the HCT116 spheroids, likely as a result of stronger actomyosin contractility induced by a 10-fold higher expression of myosin II, which led to a higher propensity of the stiffer SUM149PT emboli to bulge into the matrix of the soft vessel model. The higher expression of myosin II in the SUM149PT cells was also likely why blebbistatin—at the low, solubility-limited concentrations used—did not successfully allow growth of the SUM149PT emboli to be constrained by the soft vessel model as it was for the blebbistatin-treated HCT116 emboli. Considerably increased relaxation of the ROCK inhibitor-treated SUM149PT spheroids, such that the mechanical mismatch with the soft agarose was shifted from less than one for the elastic moduli to greater than one for the relaxed moduli, may explain their constrained growth in the soft vessel model. Finally, the prominent effect of the ROCK inhibitor on the SUM149PT emboli was not realized in the HCT116 emboli, which may be due to known cell-line dependent effects of the drug [75].

Here we demonstrated that the growth pattern of emboli could be controlled by the mismatch in the mechanical properties between a vessel-like constraint and a tumor embolus. Previous studies have shown that increasing the stiffness of a full hydrogel constraint suppresses tumor growth [19,21] and cancer cell proliferation [52]. However, in our study, increasing the constraint in the vessel-like confinement favors tumor growth. Surprisingly, we showed myosin II inhibitor-induced increases in compliance and relaxation of tumor emboli elicited confined extension of their growth in the *soft* vessel model. Such elongation may increase tumor burden due to the increased volume of tumor within the diffusion distance of nutrients and oxygen. We modeled our channel size on lymphatic vessels, where tumor emboli are often observed; however, beyond vasculature, this cylindrical geometry also applies to the mammary duct, where the majority of mammary carcinomas initiate [76,77]. Based on the results of this study, the effects of the cylindrical geometry of the mammary duct on the progression of mammary carcinomas may also be important.

## ACKNOWLEDGEMENTS

This work was supported by the National Science Foundation [NSF CAREER 1846888 to K.L.M.] and by Rensselaer Polytechnic Institute Start-Up Funds [Kristen Mills]. Likewise, we thank Elizabeth Capogna and Benjamin Liddle of Dr. Eric Ledet’s lab at RPI for assistance with printing the 3D templates, Dr. Jason Herschkowitz of the University at Albany for providing the SUM149PT cell line, and the National Science Foundation [NSF MRI 1725984] for the Leica SP8 microscope.

