## Supplemental for "Growth of tumor emboli within a vessel model reveals dependence on the magnitude of mechanical constraint"

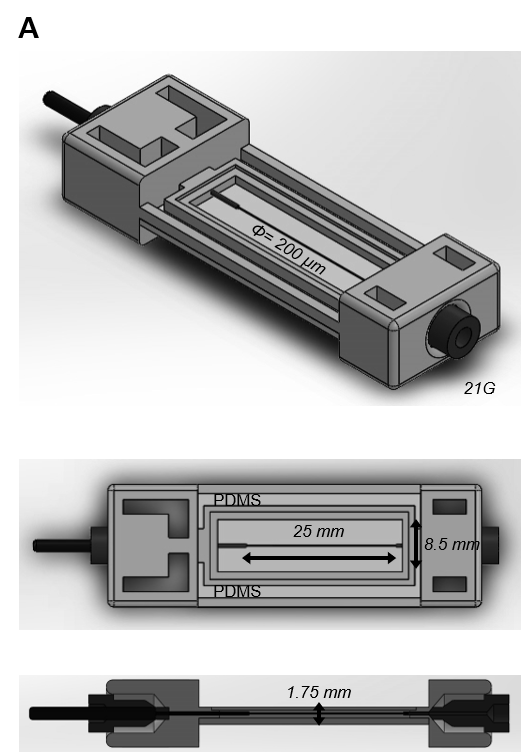

**Figure S1. Longer template for manufacturing microchannels for stiff vessel models.** Multiple views of the 3D printed template used to manufacture the stiff vessel models.

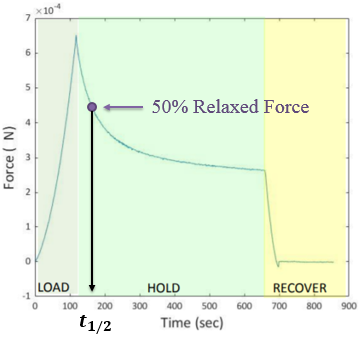

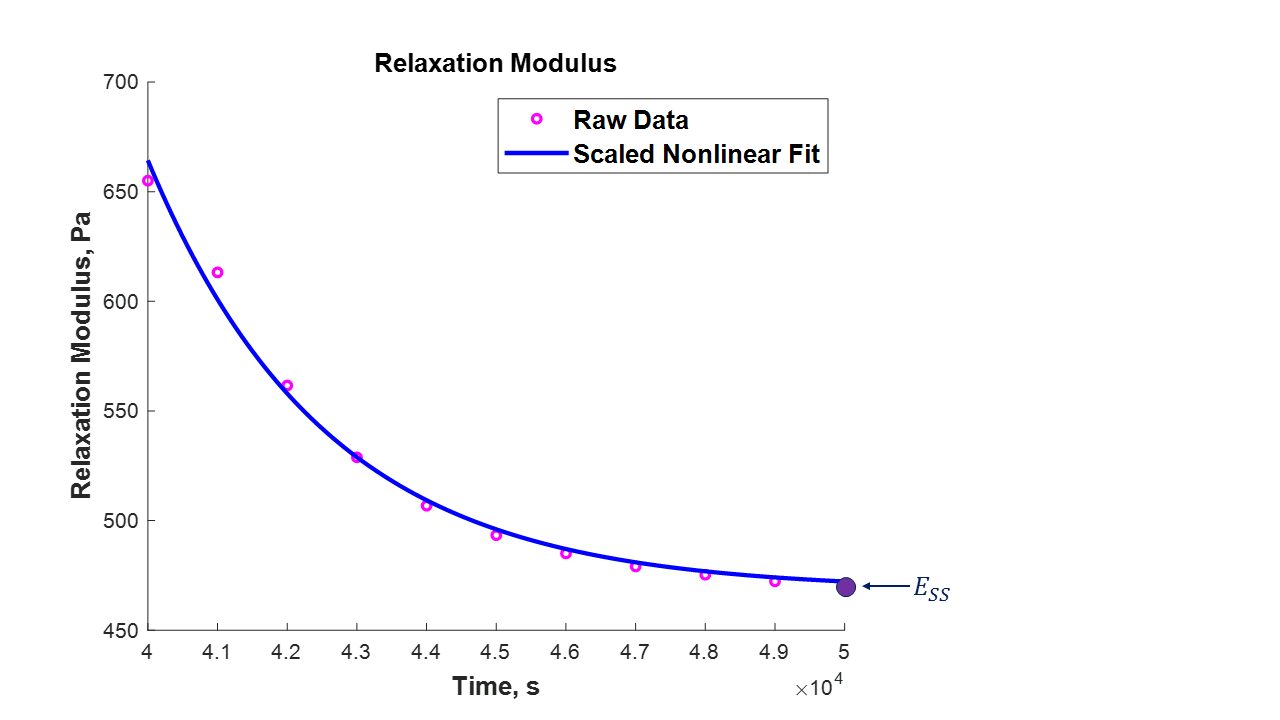

**Figure S2: Representative graphs of output and analysis from indentation and parallel plate compression experiments.** Left) Example curve showing load response of material during three test phases: indentation, hold, and retraction. Elastic indentation modulus is found from the indentation phase, and stress-relaxation properties are characterized from the hold phase. Here, half-time relaxation ($t_{1/2}$) is indicated on the horizontal axis as the amount of time required to reach half of the total force that is relaxed within the sample. Right) Example curve for time-dependent relaxation modulus (E_rel_). Here, steady state modulus (E_ss_) is shown as the final value of relaxation modulus during the hold phase, at which point the material has reached a new equilibrium.

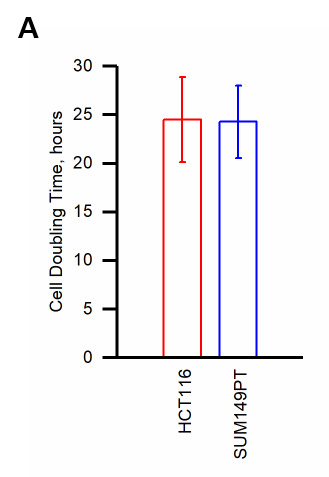

**Figure S3. Doubling time of HCT116 and SUM149P.** Graph of HCT116 and SUM149PT cell doubling time observed over three independent measurements.

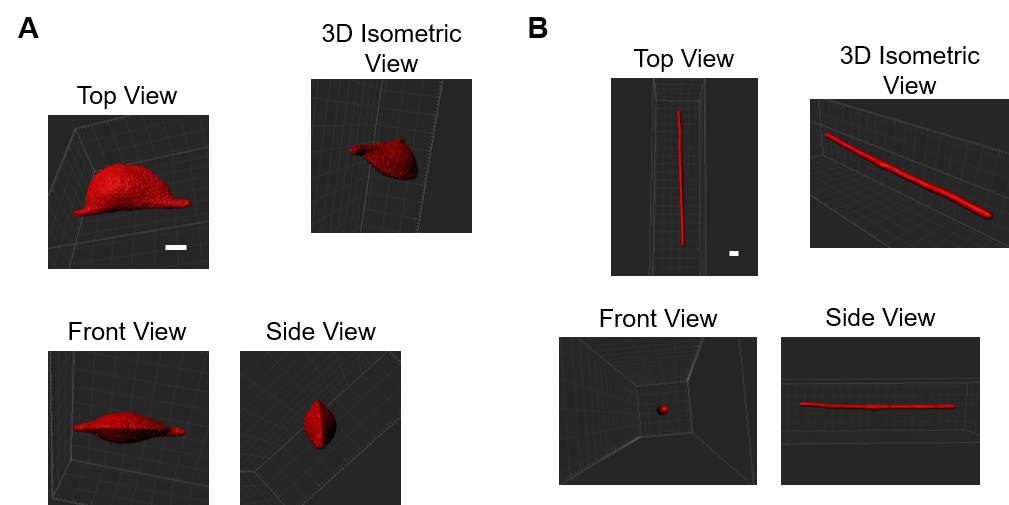

**Figure S4. MicroCT images of HCT116 emboli grown under vessel-like constraints of different stiffness**. A) Multiple views of HCT116 emboli grown under a soft vessel-like constraint after 20 days. B) Multiple views of HCT116 emboli grown under a stiff vessel-like constraint after 20 days. Scale bars are 500 µm.
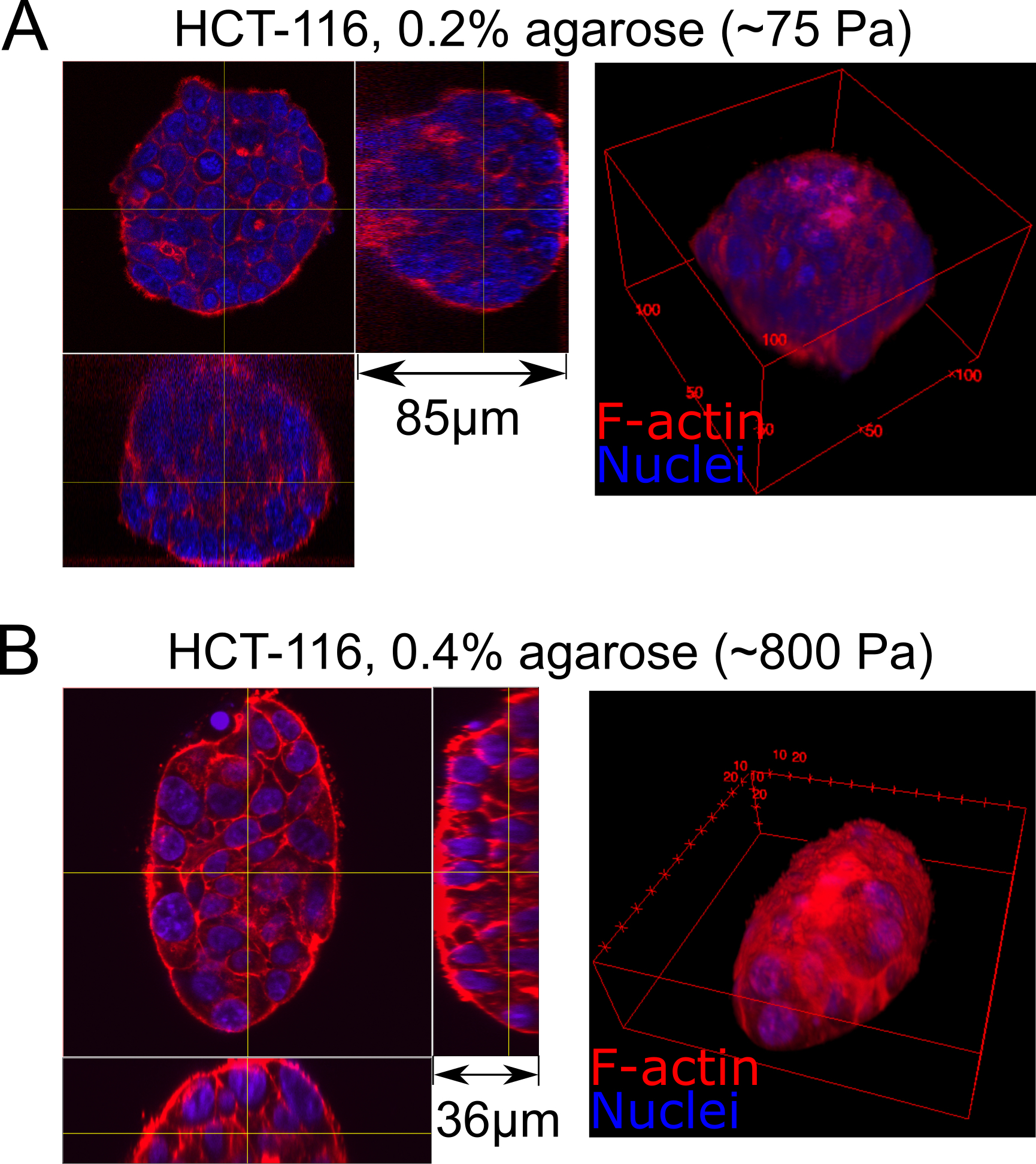

**Figure S5. An actin shell develops around tumor cell aggregates embedded in a bioinert matrix, the thickness of which is dependent on the mechanical mismatch between cell aggregate and matrix stiffness.** HCT116 cell spheroids have been measured in our study to have elastic moduli of *E_emb, HCT_* = 117 ± 31 Pa. A) When embedded and grown in 0.2% agarose with an elastic modulus of ~75 Pa—less than the stiffness of the cell spheroid—the actin layer surrounding the aggregate is *much thinner* than B) when the aggregate is embedded and grown in 0.4% agarose, which has an elastic modulus of ~800 Pa, or about 7 times greater than the cell spheroid. This excessive deposition of actin forming a shell around the aggregate may increase its apparent stiffness.

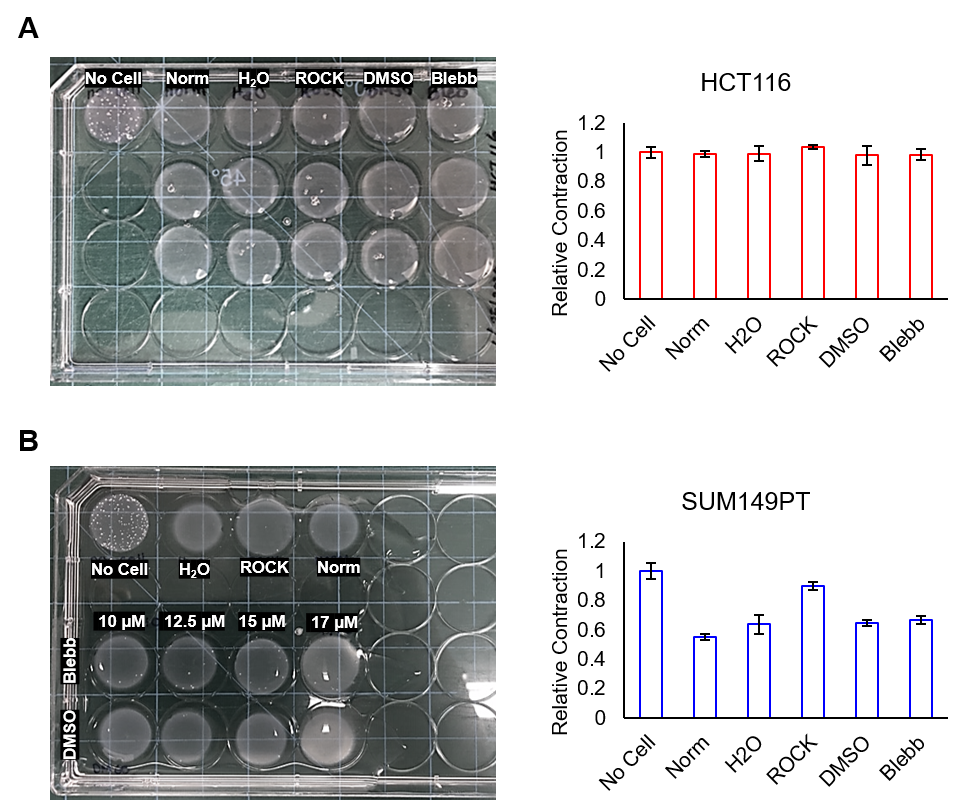

**Figure S6. Contractility measurements of HCT116 and SUM149PT cells**. A) Representative image of collagen gels containing HCT116 cells with a drug, with the solvent of a drug, or without a drug after 24 hours (left) and the measurements of relative contraction compared to a no-cell gel (right). B) Similar to (A), but for SUM149PT.

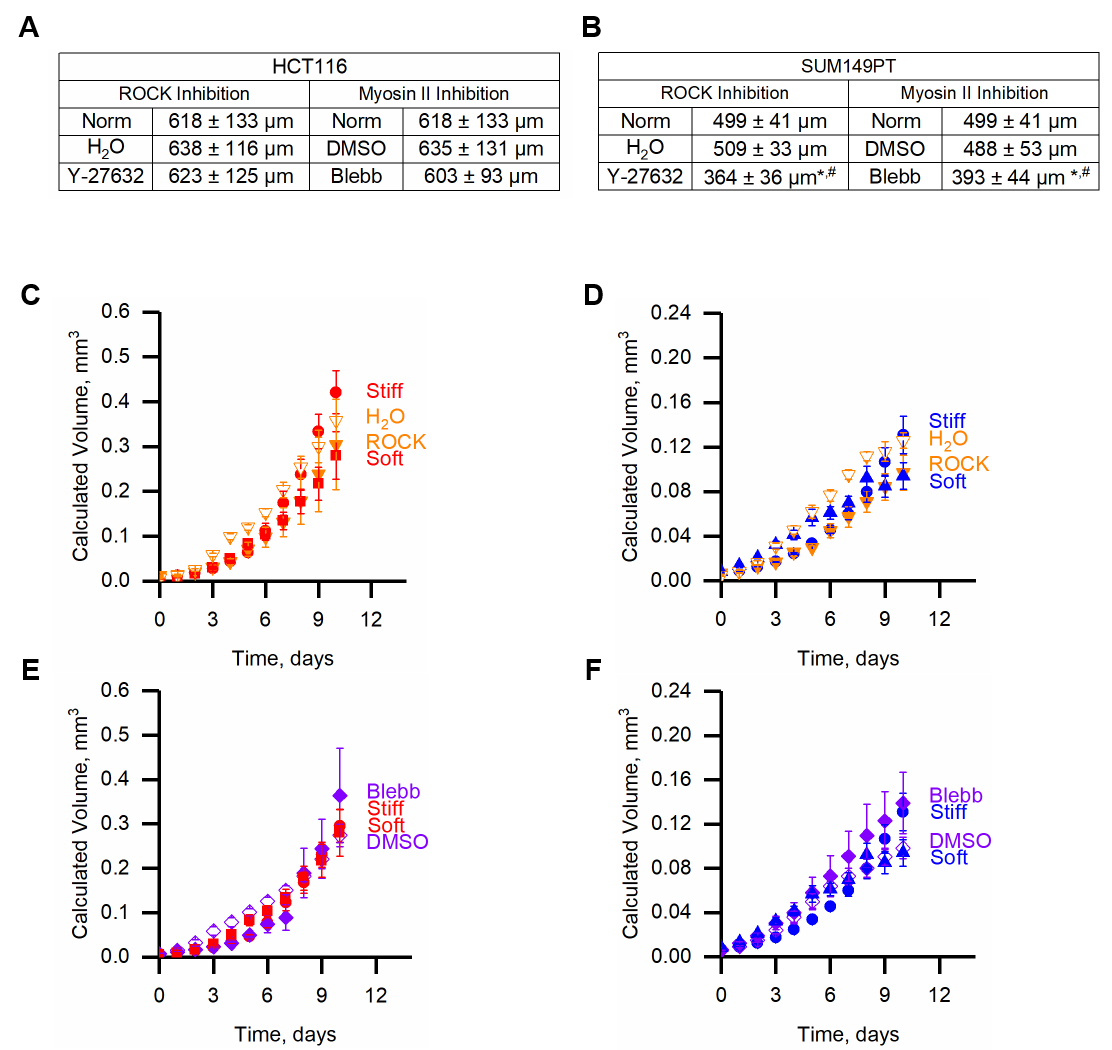

**Figure S7. Growth measurements of HCT116 and SUM149PT spheroids and emboli with the inclusion of drugs.** A) Diameters of HCT116 spheroids treated with drugs, with the drug solvent, or without drugs. B) Same as (A), but for SUM149PT spheroids. * denotes p < 0.05 compared to Norm. # denotes p < 0.05 compared to solvent control. C & E) Calculated volumes of HCT116 emboli treated with drugs, with the drug solvent, or without drugs. D & F) Same as (C & E), but for SUM149PT.

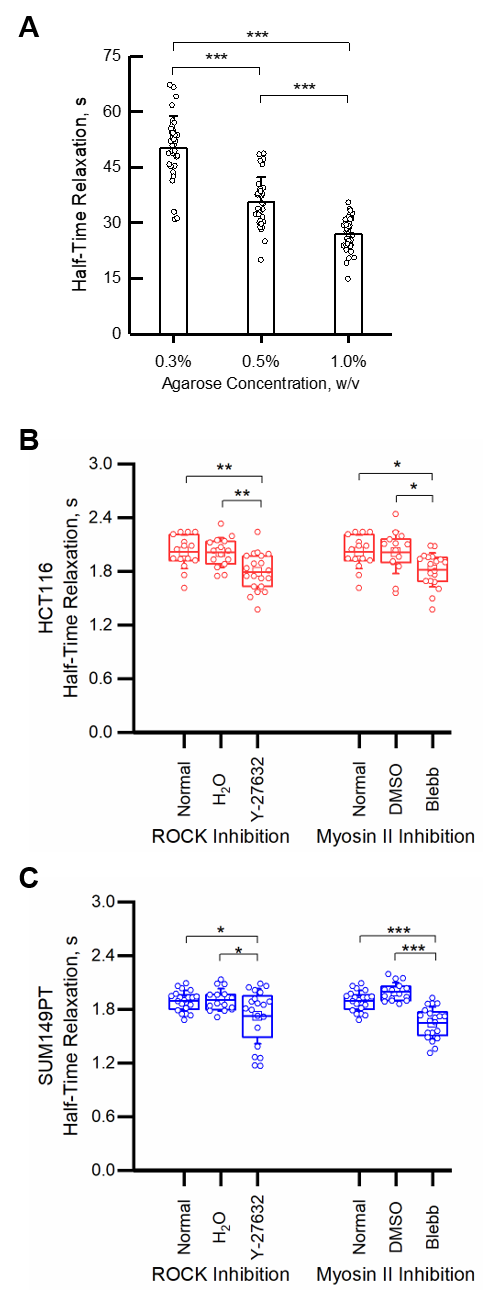

**Figure S8. Relaxation halftimes of agarose and tumor spheroids.** A) Measured half-time relaxation of agarose at different concentrations. B) Measured half-time relaxation of HCT116 with drug treatment, with drug solvent, or without drugs. C) Same as (B) but for SUM149PT spheroids. *, **, and *** denote p < 0.05, p < 0.01, and p < 0.001, respectively.
